# The 40 Hz flickering light restores synaptic plasticity and mitochondrial phenotype in experimental model of Alzheimer’s disease

**DOI:** 10.1101/2024.05.12.593775

**Authors:** Amir Barzegar behrooz, Mohamad-Reza Aghanoori, Fariba Khodagholi, Maryam Nazari, hamid Latifi, Fatemeh vosoghian, Mojdeh Anjomani, Jaber Lotfi, Abolhassan Ahmadiani, Afsaneh Eliassi, Fatemeh Nabavizadeh, Saeid Ghavami, Elham soleimani, Javad Fahanik-babaei

## Abstract

Alzheimer’s disease (AD) is the most prevalent form of dementia and a public health priority. The causes of AD are not completely understood. Pathogenetic factors including mitochondrial dysfunction, oxidative stress, reduced energy status, and compromised ion channels contribute to the onset and progression of the disease. Flickering light therapy in experimental and clinical AD has shown promising outcomes. However, the mechanisms behind the effect of flickering light at the molecular and cellular level has not yet been fully investigated. In this study, we established streptozotocin (STZ)-induced AD models by intracerebroventricular (ICV) injection of STZ in Wistar rats and monitored their memory decline. Sham and AD rats were either exposed or not exposed to 40 Hz flickering light for 7 consecutive days after 7 days of STZ injection. Memory and cognition-related behavioral analysis, pathological, electrophysiological, and biochemical assessment of the brain tissue, and mitochondrial function assays were conducted after the treatment. Cognitive and memory impairment, examined by Morris water maze (MWM), novel object recognition (NOR), and passive avoidance (PA) test, was observed in the STZ-induced AD rats and light treatment improved these behaviors. STZ injection led to significant accumulation of reactive oxygen species (ROS) and amyloid beta (Aβ), decreased serotonin and dopamine levels, and mitochondrial respiration. The 40 Hz flickering light reversed all these parameters in the light treatment group. The synaptic plasticity of STZ-induced AD rats was severely affected, but flickering light prevented the loss of synaptic plasticity and activity in the light-treated AD rats. Additionally, flickering 40 Hz white light elevated the levels of mitochondrial metabolites and the current and possible opening of the mitochondrial calcium-sensitive potassium (mitoBK_Ca_) channel which were significantly downregulated in AD rat neurons. The 40 Hz flickering light restored mitochondrial function and synaptic plasticity of neurons in AD rats and improved the cognition of animals; therefore, it can be a promising strategy to reduce AD progression.

## Introduction

Alzheimer’s disease (AD) is a progressive neurological disease. It causes cognitive decline, behavioral abnormalities, memory loss, and intellectual deterioration [1, 2]. Despite considerable efforts, pharmaceutical disease-modifying therapies for AD have not yet achieved widespread success. Most of these attempts are based on the amyloid hypothesis, which states that the disease starts with an accumulation of amyloid and then more non-specific proteins, such as total and phosphorylated tau, which cause brain damage and cognitive decline. It also initiates a chain reaction of generalized neurotoxicity events, including hyperphosphorylation, oxidative stress, local inflammation, and an imbalance between neurotransmitters such as dopamine, serotonin, and acetylcholine [3, 4]. This could result in a reduction in synaptic transmission [5], which would then impede synaptic plasticity, long-term potential (LTP), or long-term depression (LTD) [6], particularly in memory-related areas, including the hippocampus area [7].

Photobiomodulation therapy (PBMT) is a nondrug, noninvasive, and nonthermal method that uses light sources, including LASERS, and LEDs, for multiple therapeutic purposes, such as enhancing synaptic function in neurons [8–10]. Low-level LED as a therapeutic technique has been described based on sensory (visual, nasal, and/or transcranial) characteristics, which in particular, stimulation at 40 Hz reduces AD neuropathology [11–16]. Due to light flickering at 40 Hz, the ability to reverse the pathological features of AD [11, 14, 17], ischemia [18], and traumatic brain injury [19] in animal models has led to the US FDA approving a breakthrough device in 2021 for further clinical investigation [20].

Accumulation of Aβ peptides over the course of AD causes mitochondrial dysfunction, abnormal Ca^+2^ influx, synaptic loss, inflammation, and cellular death [21–25]. However, it is still unclear how Aβ causes mitochondrial structural and functional abnormalities. It is now known that PBMT improves mitochondrial function and raises the ATP content and components of respiratory chain activity, especially cytochrome c oxidase (complex IV) [26, 27]. We have shown that Aβ impairs mitochondria function in AD rat brains, and flickering 40 Hz white LEDs restores its function by raising complexes I and IV activities and mitochondria membrane potential [28, 29]. Since the mitochondria are the main sources of energy for synaptic transmission and responsible for ions/metabolites homeostasis, cell survival, and motility, it’s likely that 40 Hz light flicker produces therapeutic benefits in AD by improving mitochondrial function [30–33].

Ion channels in mitochondria are essential parts of the organelle that appear to play an important role in neuroprotection [34]. Several lines of evidence have confirmed the putative role of mitochondrial voltage- and Ca^2+^-activated K+ channel of large conductance (mitoBK_Ca_) in cell viability via the prevention of mitochondrial [Ca^+2^] overload and the reduction of reactive oxygen species (ROS) [35]. Two ATP-insensitive mitoBK_Ca_ (charybdotoxin- and iberiotoxin-sensitive) channels [36] and one ATP-sensitive mitoBK_Ca_ channel [37, 38] have recently been characterized in rat brains which are altered in Aβ and streptozotocin (STZ) toxicity models of AD and diabetes. Aβ(1-42) peptides eliminate the ATP-sensitive mitoBK_Ca_ channels, reduce conductance, and lessen the open probability of these channels in the inner membrane of the mitochondria of the neural synaptosome in AD and diabetic animal models [16, 39]. An independent study indicated that flickering 40 Hz LED could improve the activity of ATP-sensitive mitoBK_Ca_ channels in rat brains [16]. The mitoBK_Ca_ channels have not been fully investigated in STZ toxicity models of AD and the effect of 40 Hz LED on these channels remains to be determined.

In the current study, we evaluated mitochondrial metabolome as an output of the changes STZ could induce in the rat models of AD. We also studied the effect of 40 Hz LED on channel gating and permeability of mitochondrial mitoBK_Ca_ channels and if this could restore changes in the metabolomics of mitochondria.

## Materials and Methods

### Chemicals

Sucrose, D-mannitol, HEPES (4-(2-hydroxyethyl)-1-piperazineethanesulfonic acid), BSA (bovine serum albumin), EGTA (ethylene glycol-bis (β-aminoethyl ether)-N, N, N′, N′-tetraacetic acid), digitonin, nagarse, sodium bicarbonate, Trizma-HCl, potassium chloride, glibenclamide, ATP, Mannitol, sucrose, Tris–HCl, bovine serum albumin (BSA), EDTA, NADH, KCN (potassium cyanide), sodium dithionite, potassium phosphate dibasic (K2HPO4), potassium phosphatemonobasic (KH2PO4), ubiquinone-1 (coenzyme Q1), reduced cytochrome c, 2′,7′-dichlorofluorescein diacetate (DCFH-DA), and rhodamine 123 (Rh 123), mannitol, 75 mM sucrose, HEPES, EGTA, BSA, digitonin and nagarse were obtained from Sigma (Sigma-Aldrich, St. Louis, MO, USA), and *n*-decane was purchased from Merck (Merck KGaA, Dermstadt, Germany.

### Animals

Adult Wistar rats weighing 200-220g were purchased from Tehran University, Tehran, Iran. Every four rats were kept per cage under a standard condition, with a temperature of 23±1LJ°C and humidity ranging from 50-60%. The rats were also subjected to a 12-hour light/dark cycle and had free access to food and water. This study was approved by the ethics committee of the Tehran University of Medical Sciences (IR.TUMS.MSP.REC.1399.066). All the experiments were conducted based on the Guide for the Care and Use of Laboratory Animals, National Insitute of Health.

### Light Flicker Exposure Apparatus

The rats were exposed to flickering 40 Hz light in a black box with 50×50×50 cm dimensions. The LED strip was positioned inside at the top of the box and emitted white light with a visible wavelength of 425-550 nm and an average irradiance of 12 mW/cm^2^. White light pulses in a flickering 40 Hz pattern had a duration of 12.5 ms on and 12.5 ms off. LEDs’ on/off state was controlled through an AVR microcontroller circuit connected to a computer via a USB cable. Other characteristics of the method included: Duty cycle: 50%; Average power: 30 Watt; Average irradiance/average power density: 12 mW/cm; Average energy density: 10.8 J/cm^2^; Duration of exposure: 15 min/day.

### Experimental Groups

The experimental groups were assigned randomly as follows: (1) Sham: intracerebroventricular (ICV)-saline injection with no light treatment; (2) Sham+light: ICV-saline injection with light treatment; (3) AD or STZ: ICV-STZ injection with no light treatment; and (4) AD+light or STZ+light: ICV-STZ injection with light treatment. Seven days following ICV-STZ/saline injections, the rats underwent daily exposure to 40 Hz flickering white light for 15 minutes over the 7 days of exposure. Figure 1 illustrates the experimental design (Figure 1).

**Figure 1.**
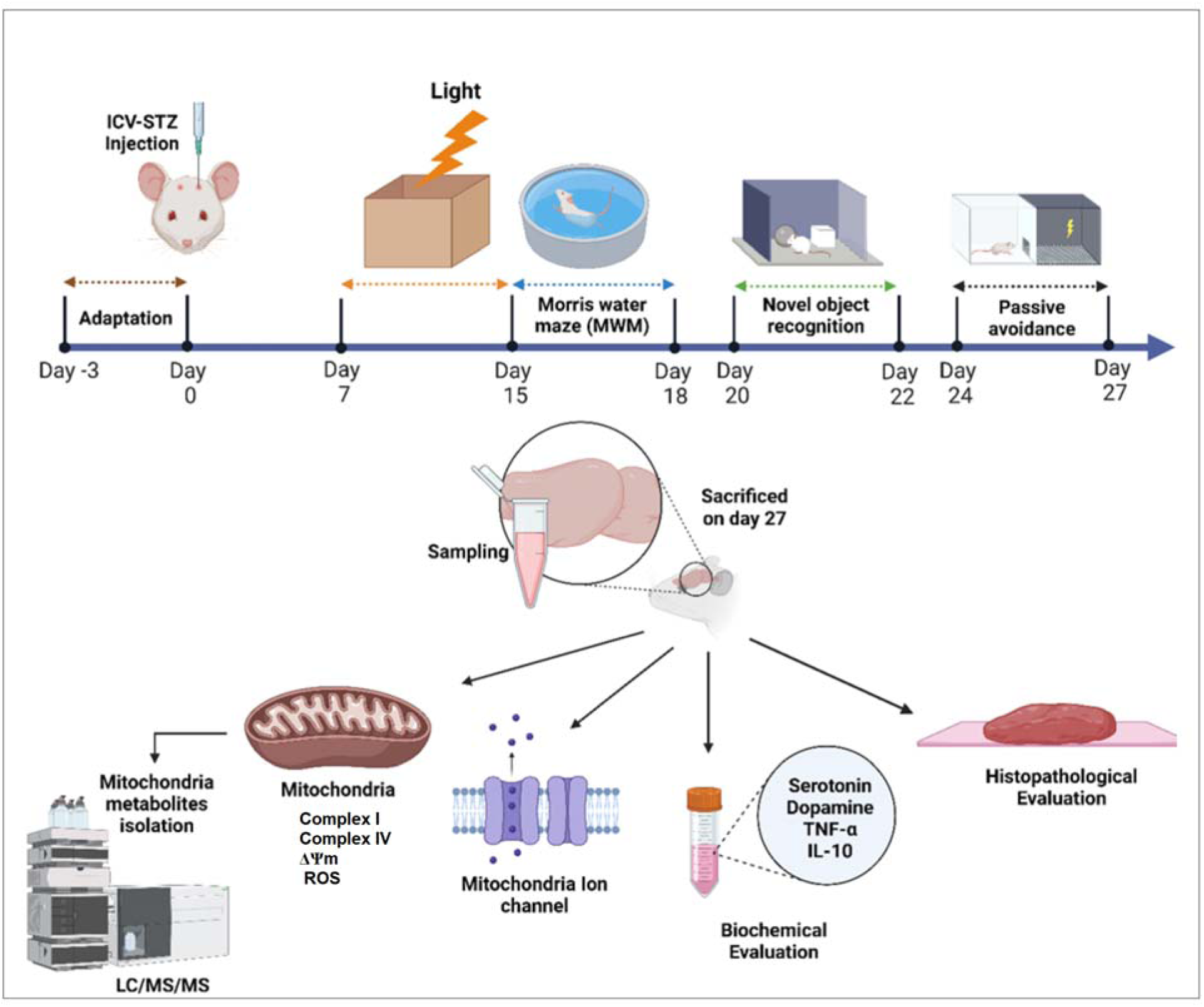
Schematic diagram of the study’s experimental design.

### Morris Water Maze

The Morris water maze (MWM) is one of the most widely used tasks to assess spatial learning and memory in rodents [40]. The task consisted of two stages: the acquisition phase for spatial learning assessment and the probe test for retention retrieval of memory. MWM was performed on our experimental rats for four consecutive days. The acquisition phase comprised one session with four trials (with 30s intervals) lasting 90s for three consecutive days. A 60s probe test with a removed platform was given 24 hours after the completion of the last session. During each trial, the path of the animals’ swimming was automatically recorded using a video camera-based system (EthoVision, Noldus, Version 14). The latency time to find the hidden platform (escape latency) was the parameter analyzed in the acquisition phase, and the latency of the first entry to the target quadrant, the number of times crossed to the platform, and time spent in the target quadrant were the parameters analyzed for probe test. One rat was deemed a ‘poor performer’ in all groups and excluded from the study.

### Novel Object Recognition Task

The Novel Object Recognition (NOR) test assesses rodents’ cognitive ability by measuring their natural tendency to explore a new object more than a familiar one [41]. The test was performed over three days beginning with a habituation period in an empty arena (60×60×50 cm) on day 1, exposure to two similar objects on day 2, and exposure to a novel object with different colors and shapes on day 3. The animals’ exploration of the objects was determined by the distance of their nose being within 2 cm of the object which was recorded using a video camera-based system (EthoVision, Noldus,Version 14). Time exploring a novel object than a familiar object and the Recognition Index (RI) was calculated based on the time spent investigating the novel object (T_N_) relative to the total object investigation (T_F_+T_N_).

### Passive Avoidance

A passive avoidance apparatus was used to assess memory retention deficits. The apparatus consisted of a wooden box divided by a guillotine door into light and dark compartments, with a stainless-steel shock floor grid in the dark section [38]. Once the rat entered the dark compartment, the door was closed, and after 30 seconds, the animal was returned to its cage. The first day consisted of three trials with 30-minute intervals, two for adaptation and the last for training. During the adaptation trial, the rats were placed in the light compartment, and their latency to enter the dark room was measured to ensure that all rats entered the dark side in under 20s. In the training phase, the rats were placed in the light chamber and allowed to enter the dark compartment. As soon as they entered, the guillotine door closed, and an electric shock (50Hz square wave, 1mA for 1.5s) was delivered to the floor. The time it took for the animal to enter the dark compartment with all four paws was recorded as step-through latency (STL). The next day, the retention latency time was measured similarly but without delivering a shock. Total time in the dark compartment (TDC) and STL were measured as memory retention evaluation, and the cutoff time was about 600s. The experiment was conducted under standard conditions between 10.00 a.m. and 2.00 p.m.

### Tissue Collection for Biochemical Tests

After behavioral tests, rats were decapitated under ketamine (150 mg/ kg) anesthesia. Rat hippocampus tissue (n = 5) was dissected and homogenized in cold RIPA buffer (pH 7.4). After centrifugation (∼10,000 g, 4°C, 15 min), the supernatant was used for biochemical tests. The Bradford assay was used to determine the total protein concentration.

### Serotonin and Dopamine Assays

Dopamine and serotonin quantities in the hippocampus tissue were assessed using the ELISA kits (Cat#: RK00642 and Cat#: TN2437, Zellbio, Germany) according to the kit instructions.

### Histological Evaluation using Congo Red Stain

Animals were anesthetized and transcardially perfused with heparinized normal saline and then 4% paraformaldehyde in 0.1 M phosphate buffer (PBS, pH 7.4). The brains were removed, and processed for sectioning and Congo red staining according to previously published protocols [42]. In the end, the hippocampus’ cornu ammonis (CA) and dentate gyrus (DG) regions on both hemispheres were subjected to neuronal Aβ plaque counting.

### Isolation of Mitochondria

The Navarro et al. protocol was utilized to isolate mitochondria [43]. Briefly, the brains were homogenized in mitochondrial isolation buffer containing 230 mM mannitol, 1.0 mM EDTA, 70 mM sucrose, and 10 mM Tris–HCl at a pH of 7.40. The homogenized brains were centrifuged at 700g for 10min, the supernatant was separated and centrifuged at 8000g for 10 to obtain mitochondria pellets. The mitochondrial samples were stored at −80°C for subsequent biochemical analysis. The mitochondria protein content was determined (0.7 mg/ml) using a Bradford assay [34].

### Mitochondrial Inner Membrane Isolation

Brain mitochondria were obtained 28 days after ICV-STZ injection using a modified version of the protocol previously published [44–46]. Briefly, dissected brains were homogenized in an MSE-nagarse solution containing 225 mM mannitol, 75 mM sucrose, 5 mM HEPES, 1 mM EGTA, 1 mg/ml BSA at pH 7.4, and 0.05% nagarse. The homogenized solution was centrifuged at 2000g for 4min, and the supernatant was centrifuged at 12000g for 9min to obtain a pellet. MSE-digitonin (0.02%) solution was used to dissolve the pellet, then centrifuged at 12000g for 11min to obtain the final pellet in cold MSE solution. The pellet was further processed according to the established protocol by Da Cruz et al. [46] to isolate and purify the inner mitochondrial membrane in the form of vesicles. To begin with, the acquired mitochondria were suspended in H_2_O at a concentration of 5 mg/ml and stirred for 20 minutes. This was followed by homogenization of the suspension 20 times using a glass homogenizer. The resulting mixture was then centrifuged twice at 12,000 g for 5 minutes and the pellet (mitoplasts) was treated with 0.1 M Na_2_C_O3_ (pH 11.5) for 20 minutes before being centrifuged again at 100,000 g for 30 minutes. Ultimately, the inner membrane vesicles of mitochondria were stored at −80 °C until required.

### Extraction of L**□α□**Phosphatidylcholine

L-α-phosphatidylcholine (L-α-lecithin) was utilized to create a planar lipid bilayer. The Singleton and Gray’s [47] method was applied to extract L-lecithin from fresh egg yolk, and the extracted lipid’s purity was verified using thin-layer chromatography (TLC), which revealed a purity of approximately 98% compared to standard lecithin.

### Single-Channel Recordings

A Delrin partition with a 150-μm-diameter hole was used to create two chambers, namely cis and trans, which served as cytoplasmic and luminal sides, respectively, for forming planar phospholipid bilayers. Both chambers were filled with 4 ml of KCl 200 mM cis/50 mM trans, and pH was adjusted to 7.2 on both sides using a Tris-HEPES solution. A solution of L-lecithin in n-decane at a concentration of 25 mg/ml was used to paint a planar lipid bilayer, which was then mechanically fused with a vesicle by gently touching the membrane from the cis side using a stainless-steel wire. Ag/AgCl electrodes were used to monitor the voltage across the bilayer, while a BC-525D amplifier (Warner Instrument, USA) in voltage-clamp mode was applied to amplify the current. To determine the thickness of the bilayer membrane, the BC-525D amplifier induced a low frequency (1-10 Hz) triangular wave with an amplitude of 5-20 mV (peak to peak), resulting in capacitance values ranging from 200 to 300 pF. The command voltage was set to the cis electrode while the trans electrode was grounded. Single-channel recordings were filtered at 1 kHz with a four-bessel low-pass filter and digitized at a sample rate of 10 kHz. The data was stored on a personal computer using Clampex10 (Axon Instruments Inc., USA) and analyzed using pCLAMP10 (Axon Instruments Inc., USA). The single-channel conductance was measured from the slope of the current-voltage curve, while the channel open probability (Po) was determined using standard event detection algorithms in pCLAMP10. Po was calculated on segments of continuous recordings lasting 40 seconds, and the results were presented as means ± S.E.

### Field Excitatory Postsynaptic Potential (fEPSP) and Long-Term Potential

To record field excitatory post-synaptic potential (fEPSP), an extracellular bipolar stainless-steel stimulating electrode with a diameter of 0.125 mm was inserted into the medial perforant pathway (4.2 mm lateral to the lambda and -3.2 mm ventrally). In comparison, another one was placed in the DG with the maximum response (-3.8 mm posterior and 2.2 mm lateral to the bregma). Extracellular field potentials were amplified 1000 times, digitized at 10 kHz, and filtered with a differential amplifier at frequencies ranging from 0.1 Hz to 10 kHz. Biphasic square waves (with a width of 200 ms) were used as stimuli. A/D interface (Science Beam Co., Iran) was used to transmit signals to a computer, and the data was analyzed using eProbe software. The stimulation intensity was adjusted to elicit a response of 40% of the maximum response population spike (PS) and fEPSP. Before inducing LTP, we conducted stimulus-response curves using various stimulus intensity levels ranging from 100 to 1200 μA. Following a minimum of 30 minutes of consistent baseline recording, we induced LTP by administering high-frequency stimulation (HFS), which consisted of 10 sets of 10 pulses at a frequency of 400 Hz, with a 10-second interval between each set. Following the tetanic stimuli, the baseline stimulation was resumed, and the recording was carried out for at least 60 minutes. The average of five successive evoked responses was taken at intervals of 10 seconds [48].

### Paired-Pulse Response

After establishing a baseline for 30 minutes, we measured the effect of paired-pulse stimulation on depression/facilitation. The stimulation was delivered at 20, 30, 50, 70, 100 and 120 ms intervals and 40% of the maximum stimulus intensity. Ten consecutive evoked responses were averaged for each paired-pulse stimulation. Various interstimulus intervals were used to determine the fEPSP slope ratio [percentage of the second fEPSP slope/first fEPSP slope; fEPSP2/fEPSP1%] and the population spike amplitude ratio [percentage of the second population spike amplitude to the first population spike amplitude; PS2/PS1%] [48].

### Tandem Mass Spectrometry (MS/MS) Analysis

After the isolation of mitochondria according to the Navarro et al. protocol [43], the acquired mitochondria were suspended in H**_2_**O at a concentration of 5 mg/ml and stirred for 10 minutes. This was followed by homogenizing the suspension 20 times using a glass homogenizer. Using the isotope dilution mass spectrometry (MS), amino acids in butyl-ester forms in the mitochondria samples were assessed using the tandem mass spectrometry system (SHIMADZU, CLAM2040, MS/MS, JAPAN) and commercial kits [CHROMSYSTEM MS/ MS complete kits, amino acids in dried mitochondrial samples spots (DBS), CHROMSYSTEM Co, Germany]. According to the kit instructions, 10 μl of the samples were transferred to a microtube, then 200 μl of internal standard was pipetted on the sample, and vortexed for 10s. Then, centrifuged at 4°C for 10min with 10000 rpm, and 150 μl of supernatant was transferred to a microplate for 20min at 45°C to dry. Afterward, 50 μl of Acetyl chloride 1-butanol was added to the sample and was left for shaking for 1 min at 700 rpm. Samples were incubated at 65°C for 15min and dried under nitrogen gas at 45°C for 15min. Then, 50 µl of 75% acetonitrile was added to the samples. Finally, samples were shaken for 2min and introduced to the MS/MS system. The injection volume was 10 μl, the flow rate was 150 μl/min, and an analysis time of 2.074min was applied. MRM (multiple reaction monitoring) was adjusted as a scan mode [49].

### Complex I (NADH**□**CoQ Oxidoreductase) Activity Assay

To determine the activity of complex I, the method described by Spinazzi et al. [41] was used with a slight modification. Briefly, mitochondria were isolated and added to an assay medium including distilled water, 500 mM potassium phosphate buffer (pH 7.5), 10 mM KCN, 10 mM NADH, and 50 mg/ml BSA without fatty acids. The reaction was initiated by adding ubiquinone-1 (10 mM) as an electron acceptor, and the decrease in absorbance was measured at a wavelength of 340 nm for 5min with 15-second intervals. The NADH has an extinction coefficient of 6.2/mM/cm.

### Complex IV (Cytochrome c Oxidase) Activity Assay

The activity of Complex IV was determined using a previously described method [50]. In brief, the cytochrome oxidation process began after adding brain mitochondrial preparation to the reaction buffer solution. The reduction in absorbance was measured at 550 nm with 15-second intervals for 5min. The enzyme activity was expressed in nmol/min/mg of protein.

### Mitochondrial Membrane Potential Measurement

Luo and Shi’s [51] protocol was used to determine the brain’s mitochondrial membrane potential (MMP). A cationic fluorescent dye called Rhodamin 123 (Rh123) was used to estimate the potential. In brief, the mitochondrial fractions were incubated with Rh123 for 5min and then excited at a wavelength of 490 nm. The emission wavelength of 530 nm was used to detect Rh123. The mitochondrial membrane potential was measured by analyzing the fluorescence intensity using a fluorescent microplate reader.

### Reactive Oxygen Species Assay

DCFH-DA was utilized to measure ROS production, as it can easily cross membranes and transform into a highly fluorescent DCF when exposed to reactive oxygen during oxidation. The isolated mitochondrial samples (0.7 mg/ml protein) were incubated with DCFH-DA in microplate wells for 20 minutes at 25°C. The fluorescence was then measured using a Multi-Mode Microplate reader (synergy™ HTX) with an excitation wavelength of 485 nm and an emission wavelength of 528 nm [52].

### Systems Biology: Metabolites-Related Pathway Identification

To identify the biological pathways affected by the metabolites dysregulated in our experiments, we used enrichment analysis, pathway analysis, and joint pathway analysis through MetaboAnalyst 6 (https://www.metaboanalyst.ca/) [53]. Before performing the joint pathway analysis, we identified the proteins related to metabolites (amino acids) through KEGG and MetaCyc, which are integrated into the Pathview Web server (https://pathview.uncc.edu/) [54].

### Statistical Analysis

Statistical analysis was performed using Prism 8 software (Graph Pad Software, USA). In all of the groups, all experimental analyses were performed blindly. Statistical analysis was performed using two-way ANOVA followed by Bonferroni’s post hoc test to compare MWM in the acquisition phase, exploring time in novel object test and fEPSP results. In addition, one-way ANOVA followed by Bonferroni’s post hoc test to compare more than two groups. The data was presented as mean ± S.E. Statistical significance was determined by considering a P value of less than 0.05 (P < 0.05).

## Results

### Flickering 40 Hz white light improved the cognitive function of STZ-induced AD rats

To investigate the function of cognition, memory, and learning in our STZ-induced AD rats, and the effect of flickering 40 Hz white light on these animals, we performed a series of relevant behavioral tests in the following groups of animals: sham, sham+light, STZ-induced AD and STZ-induced AD+light groups.

In the MWM test, the STZ-induced AD rats showed higher latency to reach the platform during the learning phase (days 1-3) compared to the sham groups (sham and sham + light) (P<0.0001). Receiving flickering 40 Hz white light for 7 days significantly shortened the escape latency in STZ-induced AD rats (P<0.01; Figure 2A). The performance of rats in all groups progressively improved during days 1 to 3 of the training session and the difference in performance remained throughout the experiment.

**Figure 2.**
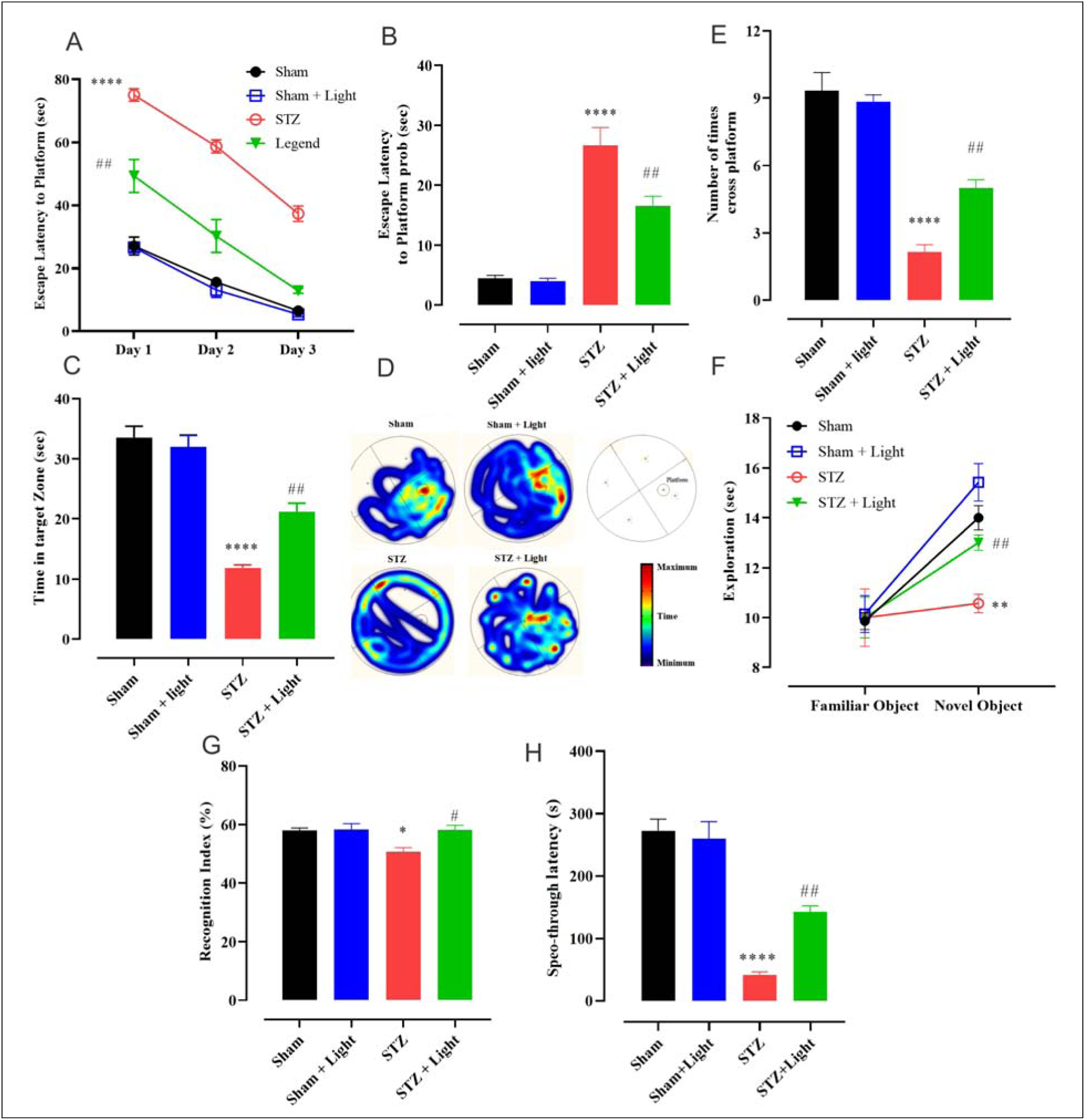
Flickering 40 Hz white light improved cognitive function of STZ-induced AD rats. MWM: Comparison of the time escape latency to reach the hidden platform in the acquisition phase for 4 days and the time escapes latency, the total time spent in the target quadrant, and several time cross-platform in probe tests in sham, sham+light, STZ-induced AD and STZ-induced AD+light groups (A-D). Escape latency to platform for the 4 rat groups during the acquisition phase (A). Bar chart for the first entry to the platform location in the 4 rat groupd (B), the total time spent in the target quadrant (C) (A comparison of the average heatmap of MWM trials between groups. The heatmap scale bar indicates the time in seconds heatmap of time spent in the target quadrant (D)), and number of time crossed the platform (E) all examined in the probe test. Novel object: Comparison of time exploring a novel object than a familiar object (F) and recognition index in all the four animal groups (G). Passive avoidance: The effect of light treatment on passive avoidance learning acquisition. Step through latency (STL) during the retrieval test performed 1 day after passive avoidance acquisition (H). One-way ANOVA analysis followed by Bonferroni post hoc test was performed to compare mean between groups. Data are expressed as mean ± SEM (n = 7 rat/group). P values are as follows: *: P<0.0.5, **: P<0.01, ***: P<0.001, ****: P<0.0001 for the sham groups vs. AD, and #: P<0.05, ##: P<0.01 for the AD groups compared to AD+light.

A probe trial test evaluated memory retrieval twenty-four hours after the last training. There was a delay in the first entry of STZ-induced AD rats to the platform location compared to the sham groups (F_(3, 24)_ = 42.73, P <0.0001) which was partially improved by the 40 Hz white light treatment (P<0.001) (Figure 2B). In this trial test, there was also a shorter time to spend in the target quadrant and less number of times crossing the platform by the STZ-induced AD rats compared to the sham groups (P<0.0001 and P <0.0001 respectively). The STZ-induced AD rats receiving flickering 40 Hz white light displayed significant improvement in these cognition parameters compared to the non-treated STZ-induced AD rats (P <0.01 and P <0.01 respectively) (Figure 2C-2E). There was no significant difference between the sham and sham-treated groups for these parameters.

Performance on the NOR task was measured by the total exploration time and recognition index. Our results showed that STZ-induced AD rats had less tendency to spend more time exploring the novel object in comparison to the sham groups as indicated by total exploration time (P < 0.001, for each subjects) and recognition index (50.16 ± 2.81%; P <0.05). Treatment of STZ-induced AD rats using flickering 40 Hz white light improved their recognition index (P<0.05). This treatment restored the behavioral deficits of STZ-induced AD rats to normal rats. Total exploration and recognition index were unaffected by flickering light in sham animals (Figure 2F-2G).

Passive avoidance behavior in shuttle box instruments involves learning to avoid certain stimuli. STL significantly decreased in STZ-induced AD rats when compared to sham group rats (P <0.0001) while receiving flickering 40 Hz white light rats increased STL by 3 folds compared to non-treated STZ-induced AD rats (P <0.01). As seen in Figure 3H, STL was unaffected by flickering light in sham animals (Figure 2H). These findings indicate that STZ-induced AD rats recall abilities and cognitive functions were enhanced by flickering 40 Hz white light.

**Figure 3.**
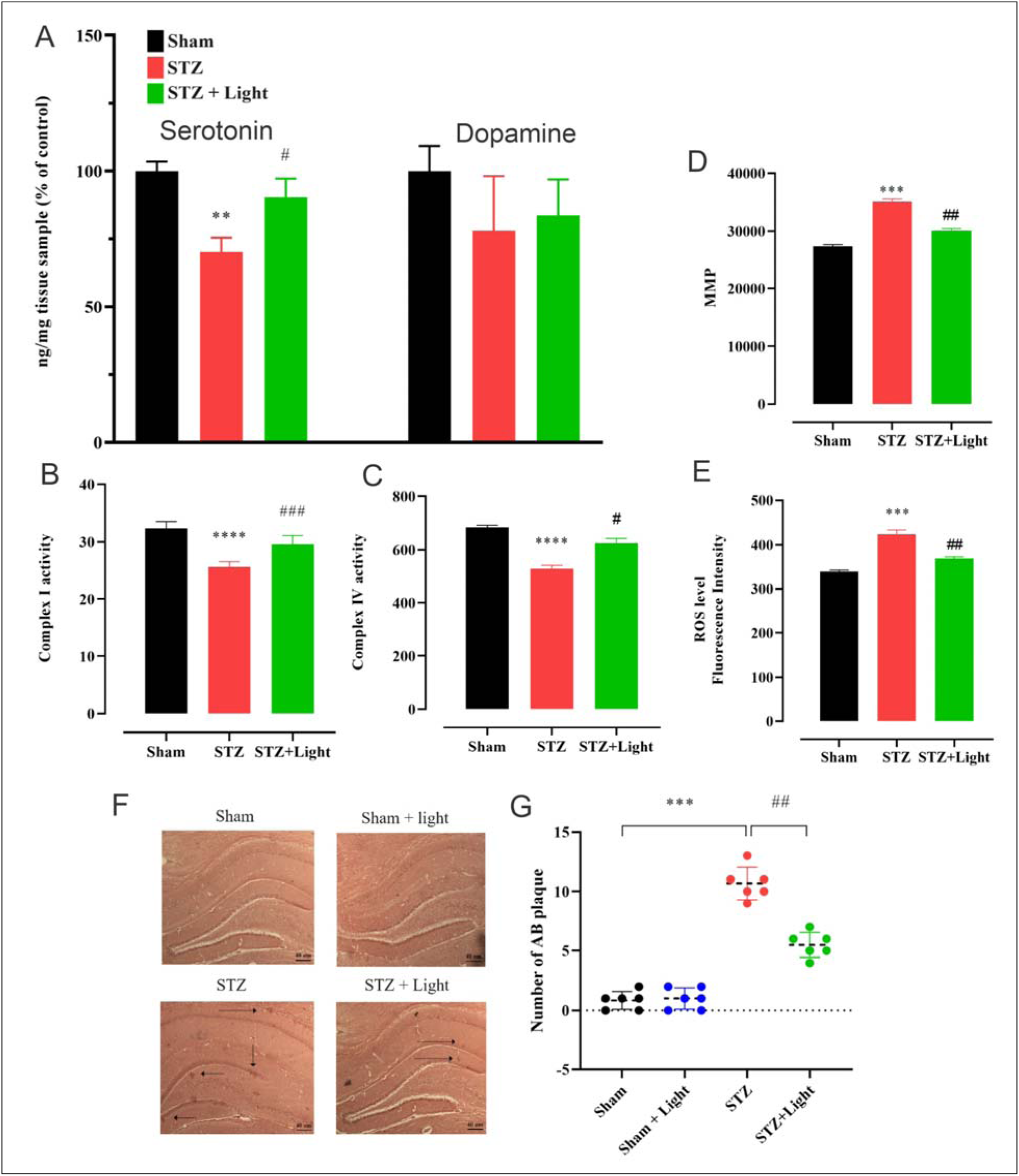
Flickering 40 Hz white light restored the deficits in serotonin levels and mitochondrial phenotype in the hippocampus of STZ-induced AD rats. Neurotransmitter (serotonin and dopamine) levels (A) and mitochondrial phenotype (B-E) were analyzed in the hippocampus tissue of sham, STZ-induced AD, and STZ-induced AD+light rats. To measure serotonin and dopamine levels, the hippocampus tissue was homogenized and subjected to ELISA assay (n = 6) (A). Mitochondria were isolated for the measurement of Complex activities, MMP, and ROS levels (B-E). In F and G, brain sections from sham, sham+light, STZ-induced AD, and STZ-induced AD+light rats were stained using Congo red, and the plaques were counted (F-G). One-way ANOVA analysis followed by the Bonferroni post hoc test was performed to compare the mean between groups. Data are expressed as mean ± SEM (n = 7 rat/group for B-E). P values are as follows: *: P<0.0.5, **: P<0.01, ***: P<0.001, ****: P<0.0001 for the sham groups vs. AD, and #: P<0.05, ##: P<0.01 for the AD groups compared to AD+light.

### Flickering 40 Hz white light restored the deficits in s**erotonin** levels **and** mitochondrial phenotype in the hippocampus of STZ-induced AD rats

At the cellular level, serotonin and dopamine levels were measured in the hippocampus tissue of the three animal groups: sham, STZ-induced AD, and STZ-induced AD+light groups (we excluded sham+light animal group from some of our analyses for simplicity since we didn’t observe any behavioral and pathophysiological differences between sham and sham+light rats). Serotonin level was significantly reduced in the hippocampus of STZ-induced AD rats vs sham group (P<0.01) while there was no significant change observed in the dopamine concentration between groups. Interestingly, the flickering 40 Hz white light treatment restored the serotonin level to normal quantities in the STZ-induced AD (P<0.05) (Figure 3A).

In the hippocampus of the three groups of animals, we further analyzed the phenotype of mitochondria reflected in the following parameters: Complexes I and IV activities, MMP, and ROS levels. Complex I and IV activities were significantly diminished in the hippocampus of STZ-induced AD rats compared to the sham group (P<0.0001). Treatment of STZ-induced AD rats using flickering 40 Hz white light for 7 days improved the activity of both complex I (P<0.001) and complex IV (P<0.05) of the electron transport chain in mitochondria when compared to the non-treated STZ-induced AD rats (Figure 3B-3C).

To further explore mitochondrial function changes in the hippocampus of these animal groups, we evaluated the fluorescence intensity of Rh123 as an indicator of MMP. There was a significant increase in the MMP of mitochondria (ΔΨm) in the hippocampus of STZ-induced AD rats compared to sham rats (P<0.001) which was driven back to normal potential when STZ-induced AD rats received flickering 40 Hz white light (P<0.01 for treated vs. non-treated AD rats)(Figure 3D). Higher ROS production was also observed in isolated mitochondria of the hippocampus tissue of STZ-induced AD rats compared to sham rats (P<0.001). This ROS overproduction was eliminated in the STZ-induced AD rats when treated with flickering 40 Hz white light over 7 days (P<0.01 for treated vs. non-treated AD rats)(Figure 3E).

### Flickering 40 Hz white light reduced A**β** plaques in the hippocampus of STZ-induced AD rats

The accumulation of Aβ in the hippocampus is a major pathological factor in the progress of AD. Congo red was used to stain brain sections derived from the following four groups of rats: sham, sham+light, STZ-induced AD, and STZ-induced AD+light groups. Clear accumulation of Aβ plaques was detected in the hippocampus of STZ-induced AD rats (P<0.001: AD vs sham). Flickering 40 Hz white light markedly reduced Aβ plaque formation in the hippocampus of STZ-induced AD rats (P<0.01: AD+light vs AD) (Figure 3F-3G).

### Flickering 40 Hz white light recovered the deficits in electrophysiological properties of hippocampal neurons in STZ-induced AD rats

To further investigate the long-term effects of flickering 40 Hz white light on neuronal activity, we assessed basal synaptic transmission and plasticity in the performant pathway of the hippocampal one month after STZ induction of AD. Basal synaptic transmission was studied by analyzing stimulus-response curves and short- and long-term potentiation by assessing paired-pulse facilitation (PPF) and LTP. The PS amplitude and the slope of the excitatory postsynaptic potential of a representative sample of the field potential and paired-pulse were recorded in lateral perforant pathway-dentate gyrus synapses of a sham, STZ-induced AD and STZ-induced AD+light groups. The acquired input/output (I/O) curve from recorded 60s periods revealed that PS amplitude increases with increasing current intensity from 100 to 1000μA. The acquired I/O measures revealed lower amplitude for the STZ-induced AD rats compared to the sham rats and flickering 40 Hz white light improved the I/O amplitude in the hippocampal neurons in STZ-induced AD rats, although these alterations were not significant (Figure 4A). After 60min of LTP induction, the dentate gyrus of STZ-induced AD rats exhibited significantly lower fEPSP compared to the sham rats (P<0.01) which were fully recovered by the flickering 40 Hz white light treatment in the light-treated AD group (Figure 4B).

**Figure 4.**
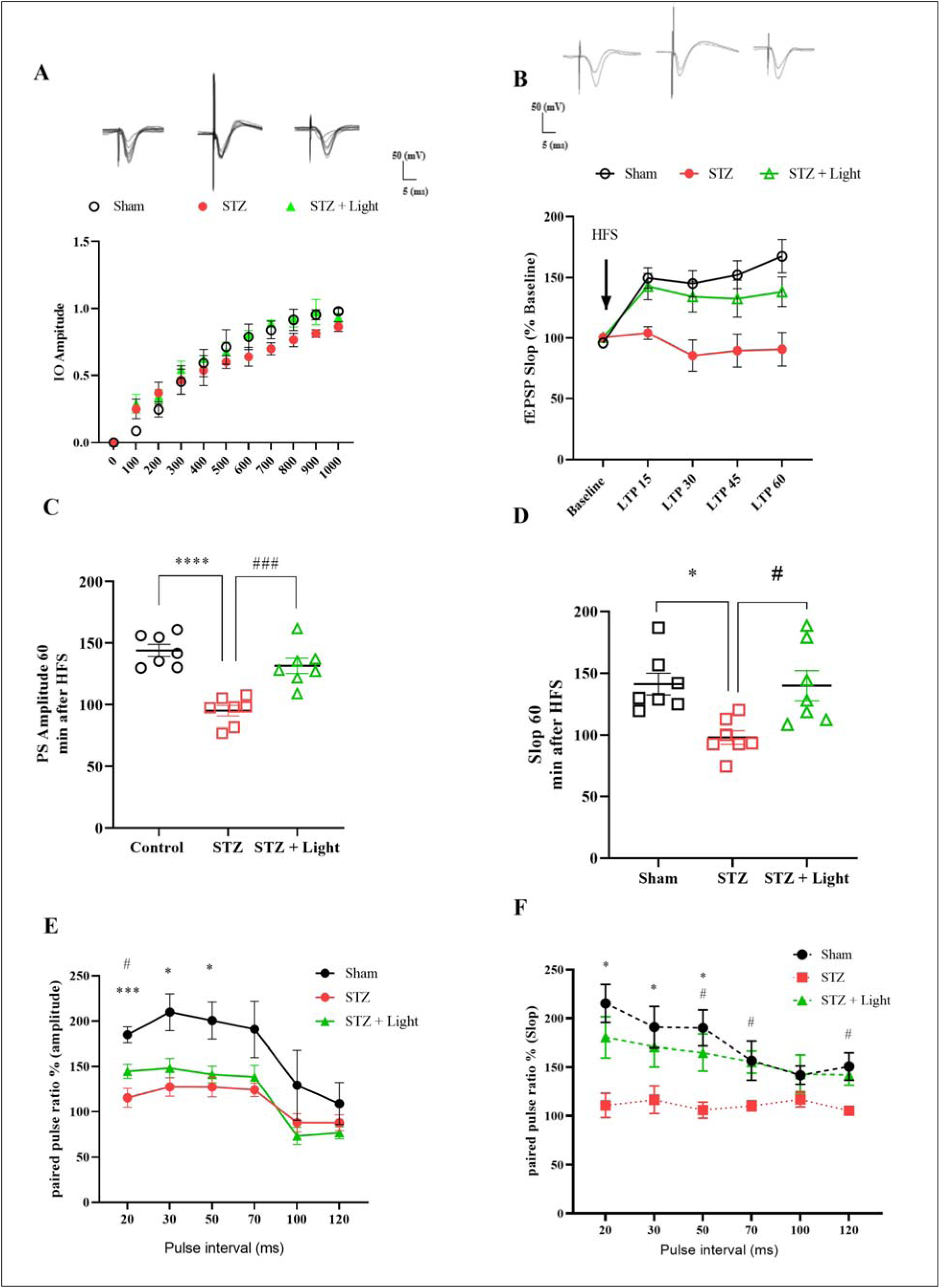
Flickering 40 Hz white light recovered the deficits in electrophysiological properties of hippocampal neurons in STZ-induced AD rats. The acquired input/output (I/O) curve from a 60s recording and single traces of the DG neurons’ reaction to numerous stimulus intensities (100 to 1000 mA) (A). The effect of flickering 40 Hz white light on LTP induction and maintenance in the dentate gyrus up to 60 min after high-frequency stimulation (HFS) (B). Data were normalized against the baseline period (-30 to +60 min), and the single traces were recorded before/after HFS. fEPSP slope change (%) and time following HFS (B). The PS amplitude and slop mean were recorded at 0 to 60min after applying HFS (n=7) (C-D). In short intervals between two strictly paired stimuli, the amplitude of PS and fEPSP slope were prompted by a paired pulse in all groups (E-F). One-way or Two-way ANOVA analyses followed by the Bonferroni post hoc test were performed to compare the mean between groups. Data are expressed as mean ± SEM (n = 7 rat/group for B-E). P values are as follows: *: P<0.0.5, **: P<0.01, ***: P<0.001, ****: P<0.0001 for the sham groups vs. AD, and #: P<0.05, ##: P<0.01, ###: P<0.001 for the AD groups compared to AD+light.

High-frequency stimulation (HFS: 400 Hz) of lateral perforant pathway-dentate gyrus produced high PS amplitude and a long-lasting synaptic potentiation which were both depressed in STZ-induced AD rats (P<0.0001 and P<0.05, respectively). Intriguingly, the flickering 40 Hz white light treatment normalized both parameters in the light-treated AD rats (Figure 4C-4D).

In Figures 4E and 4F, short intervals were given between two strictly paired stimuli, and the PS amplitude and fEPSP slop were analyzed. Repeated measures for acquired input/output revealed a significantly lower PS amplitude in the STZ-induced AD rats compared to sham rats (F_(1.65, 6.62)_ = 20.17, P <0.0006) which was recovered by 40 Hz light treatment at 20ms intervals (Figure 4E). Similarly, significantly lower fEPSP slop (F_(2.06, 8.26)_ = 4.37, P =0.049) was observed for the neurons in the dentate gyrus of STZ-induced AD rats. To our surprise, the flickering 40 Hz white light restored the defect in synaptic potentiation at 20, 30, and 50ms intervals of paired pulses in the hippocampal neurons of light-treated AD rats (Figure 4F). Overall, electrophysiological properties of hippocampal neurons including synaptic transmission and plasticity were defected in the STZ-induced AD rats and were recovered by flickering 40 Hz white light.

### Induction of AD caused a diminished channel gating and conductance in mitochondrial mitoBKCa channel, and 40 Hz light therapy reversed these defects

To explore if the biophysical properties of mitochondrial mitoBK_Ca_ channels are affected by STZ injection and flickering light, the mitochondrial inner membrane was extracted as vesicles, and channel activities were studied. Following the incorporation of vesicles extracted from the brain mitochondrial inner membrane, the success rate of the mitoBK_Ca_ channel was ∼45%, 36%, and 40% of recordings for sham, STZ-induced AD, and STZ-induced AD+light rats, respectively (data not shown). Figure 5A shows examples of single-channel activities prepared from the brain mitochondrial inner membrane vesicles in 200 mM KCl cis/50 mM KCl trans condition in the sham, STZ-induced AD, and STZ-induced AD+light animal groups, respectively (Figure 5A). The channel conductance analysis showed a ∼56% reduction (124± 7.2) in the mitochondrial inner membrane vesicles in STZ-induced AD rats compared to sham rats (channel conductance of 211) (Figure 5A-5C). As shown in Figure 5B-5C, we observed close to a 1.5-fold increase in unitary current amplitude in light-treated AD rats relative to the non-treated AD rats. Data have been summarized in Figure 5B-5C, where the slope of the I-V relationship estimates a channel conductance of 194± 11.5 pS in the light-treated AD group in contrast to 124 ±7.2 pS in the non-treated AD rats (Figure 5A-5C). Channel gating behavior indicated a voltage-dependent property, with increased longer silent periods at potentials lower than −40 mV. Figure 5A-5B indicates an outward current at potentials above −10 mV and a zero current potential value close to − 30 mV; the reverse potential is expected for potassium ions. This result confirmed the cationic nature of the channel.

**Figure 5.**
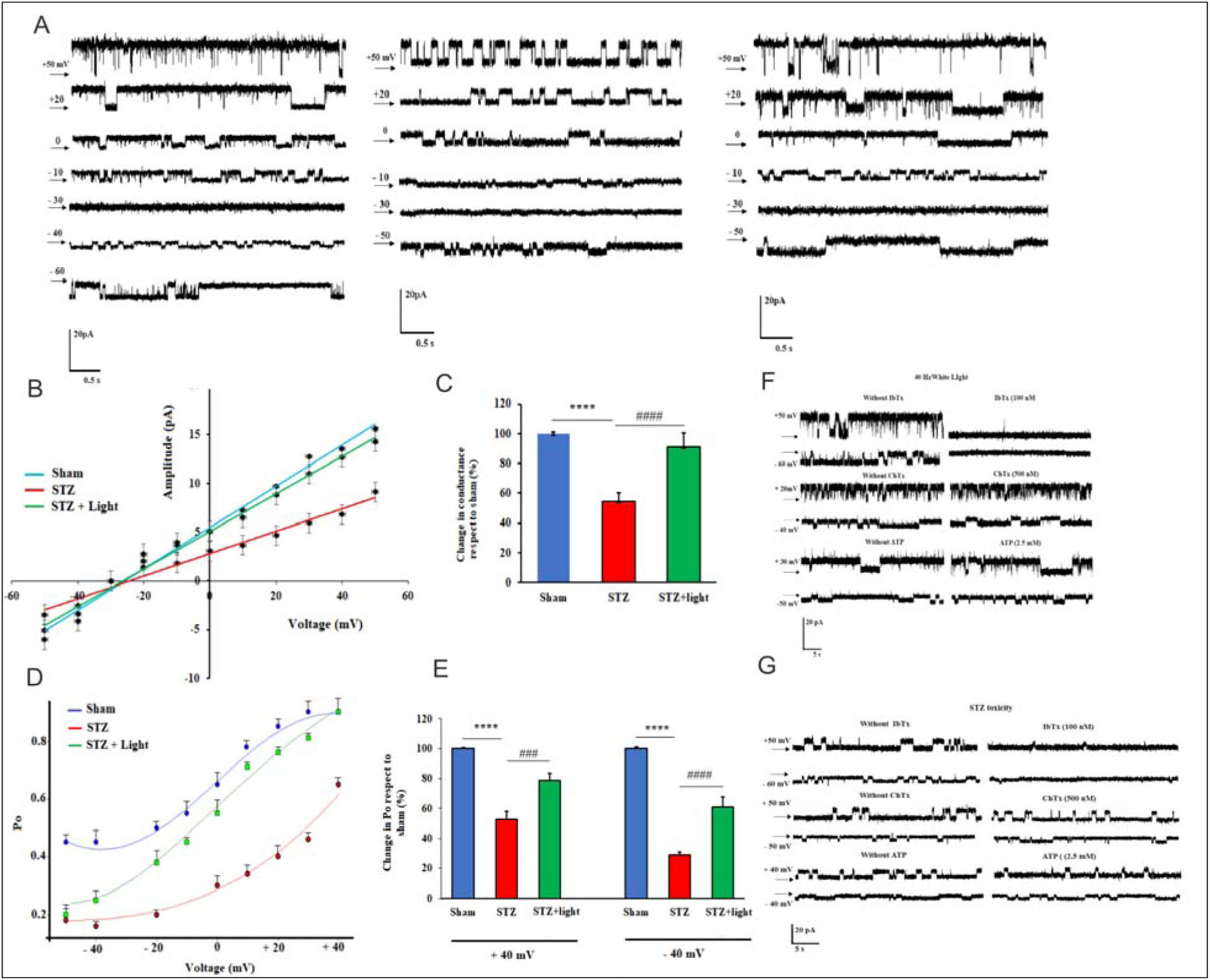
Induction of AD caused diminished channel gating and conductance in mitochondrial mitoBKCa channel, and 40 Hz light therapy reversed these defects. Single channel recording in 200/50 mM (cis/trans) KCl solution was performed using the bilayer lipid membrane. Single channel current-voltage relationship for the mitoBK_Ca_ channel and percent of changes in the channel’s conductance and open probability (Po) as a function of voltage for mitoBK_Ca_ channels was performed in animal groups (A-E). The closed state is marked by the black arrow. Each point represents the average open probability as a function of voltages from five different experiments. The effect of IbTx, ChTx, and ATP on the mitoBK_Ca_ channel activity in the STZ-induced AD (F) and STZ-induced AD+light rats (G). One-way ANOVA analyses followed by the Bonferroni post hoc test were performed to compare the mean between groups. Data are expressed as mean ± SEM (n = 5 rat/group). P values are as follows: *: P<0.0.5, **: P<0.01, ***: P<0.001, ****: P<0.0001 for the sham groups vs. AD, and #: P<0.05, ##: P<0.01, ####: P<0.0001 for the AD groups compared to AD+light.

Channel open probability (Po) values were close to 0.9 ± 0.05 at + 40 mV in sham rats. In contrast, channel Po in STZ-induced AD rats exhibited a decrease of 0.29 ± 0.02 at − 40 mV and 0.52 ± 0.04 at + 40 mV (Figure 5D-5E). Curve fitting the observed Po voltage dependence to the Boltzmann distribution indicated a voltage for half-maximal activation (V1/2) of − 67 ± 1.14 mV in the sham rats compared to 40.5 ± 4.2 mV in the STZ-induced AD rats, with an approximately equivalent gating charge zd of − 2.74 ± 0.42 and − 2.34 ± 0.43, respectively. Interestingly, flickering 40 Hz white light therapy increased the channel Po in STZ-induced AD rats from 0.16 ± 0.029 to 0.24 ± 0.05 at − 40 mV and from 0.52 ± 0.03 to 0.67 ± 0.04 at + 40 mV, respectively (Figure 5D-5E). Curve fitting the observed Po voltage dependence to the Boltzmann distribution demonstrates a V_1/2_ of − 45.25 ± 4.1 mV in the STZ-induced AD+light groups, with an equivalent gating charge zd of − 2.74 ± 0.86. These results confirm that flickering 40 Hz white light therapy improves channel activities in the STZ-induced AD rats (Figure 5D-5E).

The effects of IbTx, ChTx, and ATP were examined to verify the type of the mitoBK_Ca_ channel extracted from the rat brains of both STZ-induced AD and STZ-induced AD+light rats. Further experiments were conducted to distinguish between two different types of mitoBK_Ca_ channels including ATP-sensitive mitoBK_Ca_ channel (conductance of 565 ps and inhibited by IbTx but not ATP and ChTx) [55], and mitoBK_Ca_ channel (conductance of 211 ps and inhibited by IbTx but not ATP and ChTx [55]. A single-channel behavior was considered before and after adding each drug to the same planar lipid bilayer.

As shown in Figures 5F and 5G, channel activity was inhibited entirely after adding 100 nM IbTx at negative and positive potentials in STZ-induced AD and STZ-induced AD+light rats (n = 4 in each group). Also, no inhibition of channel activity was observed following the addition of 2.5 mM ATP (at + 40 mV and − 40 mV for STZ-induced AD, and +20 mV and -40 mV for STZ-induced AD+light. Additionally, 500 nM ChTx (at + 50 mV and − 50 mV in STZ-induced AD and + 30 mV and − 50 mV in STZ-induced AD+light rats (n = 4) did not inhibit the channel activity. These results verify that the channels obtained from rat brain preparations of STZ-induced AD and STZ-induced AD+ligh rats were the mitoBK_Ca_ channel (conductance of 211 ps and only inhibited by IbTx) (Figure 5F-5G).

### Flickering 40 Hz white light reversed the disruption in the mitochondrial metabolites quantities in STZ-induced AD rat brains

Amino acids in mitochondria, as mitochondrial metabolites, were measured using MS-MS to inspect any changes in these metabolites as a result of the modified function of mitoBK_Ca_ and mitochondrial function. There were ten amino acids with significant reduction in the STZ-induced AD rat brains compared to the sham rat brains [Tyrosine (P=0.0006), Tryptophan (P=0.0346), Alanine (P=0.0338), Leucine (P=0.0001), Methionine (P=0.0023), Proline (P=0.0172), Lysine (P=0.0076), Serine (P=0.0237), Aspartic acid (P=0.0499) and Glutamic acid (P=0.0001)]. Interestingly, flickering 40 Hz white light therapy restored the amounts of these amino acids back to normal amounts in sham rat brains (Figure 6A-6C).

**Figure 6.**
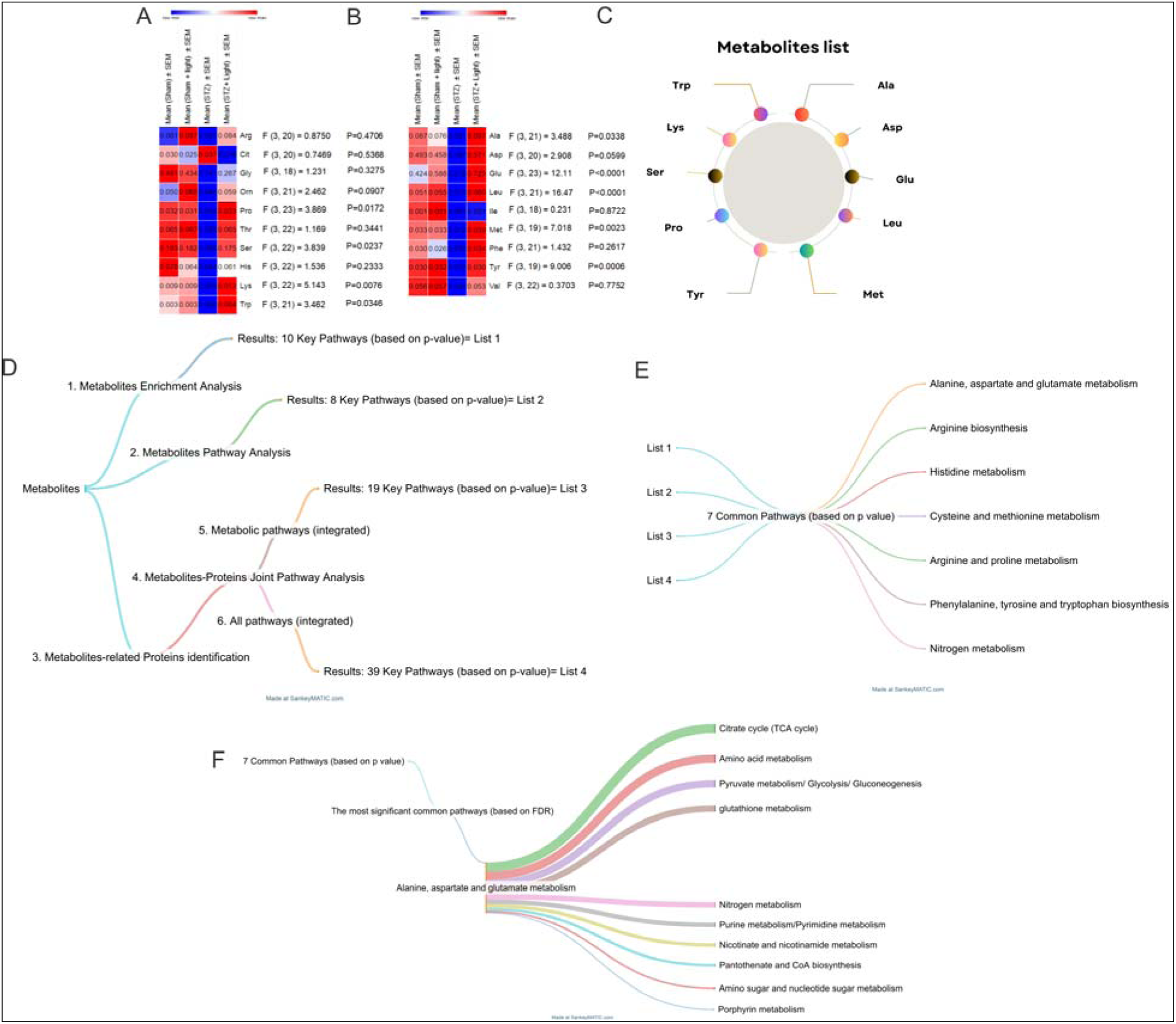
Flickering 40 Hz white light reversed the disruption in the mitochondrial amino acid quantities in STZ-induced AD rat brains. Differential levels of mitochondrial amino acids measured using MS-MS were compared in sham, sham+light, STZ-induced AD, and STZ-induced AD+light rat groups (A-B). Ten significant metabolites were downregulated in STZ-induced AD rats vs sham groups (C). Diagram of metabolite-protein joint pathway analyses (D), extracted seven common pathways (E), top 10 associated pathways affected by alanine, aspartate, and glutamine metabolism (F) are illustrated. Data are expressed as mean ± SEM (n =7±1).

### Metabolites dysregulated in STZ-induced AD rats were mostly enriched in pathways regulating the Krebs cycle (citrate cycle/TCA), carbohydrate metabolism, and amino acid metabolism in mitochondria

The integration of the results obtained from KEGG and MetaCyc, as discussed in the methods section, finally revealed a final list consisting of 154 proteins (Supplementary Table 1). By considering a p-value < 0.05, 10 specific pathways were identified after enrichment analysis (Supplementary Table 2), and 8 distinct pathways were identified after pathway analysis (Supplementary Table 3). Moreover, metabolite-protein joint pathway analysis was performed in two contexts: i. metabolic pathways (integrated) and ii. all pathways (integrated). Metabolic pathways (integrated) approaches revealed 19 distinct signaling pathways (Supplementary Table 4), whereas all pathways (integrated) determined 39 key pathways (Supplementary Table 5) (Figure 6D).

In the next step, the seven common key pathways were identified by integrating four lists. These seven pathways included Alanine, aspartate and glutamate metabolism (rno00250), Arginine biosynthesis (rno00220), Histidine metabolism (rno00340), Cysteine and methionine metabolism (rno00270), Arginine and proline metabolism (rno00330), Phenylalanine, tyrosine and tryptophan biosynthesis (rno00400), and Nitrogen metabolism (rno00910) (Figure 6E).

The most significant signaling pathway (based on FDR < 0.05) was Alanine, aspartate, and glutamate metabolism (rno00250) which involved in 10 key pathways (Figure 6F). Alanine, aspartate, and glutamate metabolism and 10 associated pathways were tightly linked to STZ-induction of AD and recovery by the flickering 40 Hz white light treatment in rats.

## Discussion

We demonstrated in the present study that flickering 40 Hz white light has beneficial effects on the brain of STZ-induced AD rats and has the potential for treating Alzheimer’s disease. We first showed that STZ induction of AD caused behavioral disorders, mitochondrial dysfunction (including its components and metabolites), plasticity impairment, neuronal death, neurotransmitter changes, and beta-amyloid plaque accumulation in rats. When flickering 40 Hz white light was used as treatment, it improved cognition, dopamine and serotonin levels, and long-term and short-term memory in AD rats. Further, we showed for the first time that flickering 40 Hz white light optimized mitochondrial function, current changes, and the possibility of opening of mitoBK_Ca_ in rat neurons.

As expected, we observed deficits, including cognitive memory deficits in the MWM, NOR, and PA tests, as previously reported in the rats subjected to ICV-STZ in AD research [56, 57]. After treatment with flickering 40 Hz white light, cognitive memory impairment induced by STZ was attenuated, indicated by appearing to decreased escape latency, increased time target and number of crosses in the prob test of MWM, increased RI in the NOR test, and increased step-through latency in passive avoidance test. Similarly, animal studies have suggested that photomodulation regulates cognitive memory [58, 59]; notably, it reduces cognitive impairments in AD rats [28, 60]. A recent study found that flickering 40 Hz white light ameliorated memory impairments in animal models of AD through inhibition of mitochondria dysfunction, oxidative stress, and suppression of apoptosis [29, 61]. These results support the idea that flickering 40 HZ white light has an anti-amnesic effect when used in an STZ-induced model of sporadic Alzheimer’s disease.

Serotonin and dopamine are monoaminergic neurotransmitters in the brain which are downregulated in AD patients and animal models [62–64]. Serotonin is the earliest neurotransmitter downregulated in the brain of AD patients which regulates the release of other neurotransmitters including dopamine [65]. Light therapy has been shown to decrease serotonin transporter binding, improve serotonin levels, and increase dopamine in human and rat [66–68]. In line with our results, dopamine levels were elevated under 40 Hz light treatment. In the dark, levels of dopamine are reduced in our body and are elevated during our light cycle of the day, as opposed to melatonin levels [16, 69, 70]. The increase we observed in the levels of both neurotransmitters might be due to elevated energy supply for synaptic activities, increased intracellular calcium levels, and light-induced neuronal activity [71, 72].

Decreased neuronal plasticity is another hallmark of neurodegenerative diseases such as AD [73]. Neuronal plasticity is directly related to greater synaptic efficacy achieved through long-term potentiation (LTP). In animals, LTP is the long-term modification in synaptic strength induced by electrical stimulation which is used in AD research studies [48, 74]. To further investigate the long-term effects of flickering 40 Hz light and bilateral injection of STZ into the hippocampus on neuronal activity, we assessed basal synaptic transmission and plasticity in perforant hippocampi one month after injection. Our observation of reduced LTP and PPF in the hippocampus of STZ-induced AD rats can be due to compromised activity of NMDA receptor and constant increase in Ca^2+^ influx caused by STZ toxicity [75]. Similar studies have shown that short-term plasticity is dependent on residual Ca^+2^ in pre-synaptic terminals, and STZ causes calcium imbalance in pre- and post-synaptic regions [76]. Flickering 40 Hz light might counteract the action of STZ on NMDA receptors and modulate Ca^2+^ levels in the cells thus preventing suppression of LTP and synaptic plasticity [77].

Our histological results indicate overproduction and formation of Aβ plaques in the hippocampus of STZ-induced AD rats. Accumulating evidence shows that Aβ is physically localized in the mitochondria and interacts with mitochondrial proteins which might explain its dysfunction [78]. Few studies have shown that Aβ in the form of oligomers can directly lead to the inhibition of the mitoBK_Ca_ channel activity which might cause mitochondrial dysfunction [21, 79]. These alterations were reported in the STZ-induced diabetic animal models [80]. In type 2 diabetic mice, insulin resistance in the periphery and the brain was associated with energy impairment [81–83]. O’Malley et al. have shown that insulin activates mitoBK_Ca_ channels in the hippocampal cultured cells and modulates spontaneous [Ca^2+^] oscillations [84]. It’s possible that insulin resistance in the brain of our STZ-induced AD rats in cooperation with Aβ plaques leads to decreased mitoBK_Ca_ channel activity, mitochondrial membrane potential, and mitochondrial dysfunction [80]. Mitochondrial BKCa channel preserves mitochondrial function and plays a critical role in neuroprotection through inhibition of mito [Ca^2+^] overload and reduction of total ROS production [35, 85]. Our finding that intact animal IbTx, but not ATP and ChTx, inhibited the channel gating behavior confirms that the AD channel corresponds to the mitoBK_Ca_ (ATP- insensitive mitoBK_Ca_ channel) according to the previous reports [36, 55].

Elevation of cytoplasmic calcium can be explained by light-induced opening of calcium ion channels, such as members of the transient receptor potential (TRP) superfamily [86]. Since nanostructured water exists in all ion channels and is sensitive to heat and light, it’s feasible that the mitoBK_Ca_ channel phenotype is also changed under light treatment. Complex IV in ETC is no exception as an ion channel and might be affected by light treatment in a similar manner [87, 88]. From a mechanistic point of view, the emitted photons in the photobiomodulation are absorbed by complex IV to facilitate the availability of electrons for the reduction of oxygen, which increases mitochondrial membrane potential, ATP levels, and ETC activity [89, 90]. Studies also have suggested a direct structural-functional relationship between the ETC and the mitoBK_Ca_ channel. It is proposed that a redox signal from the respiratory chain is transmitted to the mitoBK_Ca_ channel through cytochrome c oxidase (complex IV) [91].

Studies on mice showed that the 590nm LED light significantly increased NO synthesis in the brain through stimulation of complex IV activity in mitochondria in hypoxic conditions, and through a feedback mechanism complex IV enzyme activity was inhibited by NO [92]. Photons of 40 Hz light absorbed by complex IV can dissociate this inhibitory NO in the presence of sufficient oxygen [9]. Dissolution of NO resulted in increased MMP, higher consumption of oxygen, metabolism of glucose, and generation of more ATP. This, in turn, triggers minimal quantities of ROS followed by activation of ROS-induced signaling pathways which have cytoprotective effects in the cells [93]. On the other hand, flickering 40 Hz white light might prevent mitochondrial [Ca^2+^] overload by enhancing ETC and mitoBK_Ca_ channel activities [85]. Optimal levels of mitochondrial [Ca^2+^] prevent the overproduction of ROS, abnormally high MMP and mitochondria dysfunction [94]. This eventually promotes the viability of cells in the hippocampus and cortex, and improves cognition [26, 95]. Mitochondrial BK_Ca_ channel opening under light treatment which keeps the calcium ions in the cytoplasm, might trigger the release of the neurotransmitter glutamate from the presynaptic neuron and improve the LTP in light-receiving animals [96, 97].

In our findings regarding metabolomics analysis, ten mitochondrial amino acids were significantly decreased in STZ-induced AD rats which are classified into three categories: 1) Aromatic (Tyrosine, Tryptophan), 2) Aliphatic (Alanine, Leucine, Methionine, Proline), and 3) others (Lysine, Serine, Aspartic acid, Glutamic acid). Through different bioinformatics analyses explained in the results and methods sections, seven common pathways which had more enzymes/proteins affected by these ten amino acids were characterized. Among these, alanine, aspartate, and glutamate metabolism was the top hit with the lowest FDR and P values. In line with our findings, in a proteomic-metabolomic study of high-sugar-fat diet Wistar rats (an animal model of metabolic syndrome), 16 pathways were enriched. Three pathways were shared by metabolomics studies of serum samples from diabetic patients: 1) alanine, aspartate, and glutamate metabolism, 2) glycine, serine, and threonine metabolism, and 3) valine, leucine, and isoleucine biosynthesis [98]. Transcriptomic and metabolomic joint analysis after an immune challenge with LPS in Zebra fish, revealed enrichment of alanine, aspartate, and glutamate metabolism and aminoacyl-tRNA biosynthesis [99]. They reported Krebs cycle was one of the most affected cycles in the mitochondria and highlighted in both studies when analysis was performed by the KEGG pathway [98, 99].

The top three (out of ten) pathways mostly affected by alanine aspartate, and glutamate metabolism derived from our metabolomics-protein joint pathway analyses were TCA and glycolysis, amino acid metabolism, and glutathione metabolism. Krebs cycle and glycolysis are two key components inside and outside of the mitochondria, respectively, for energy production in the form of ATP in neurons and other cells [100–102]. In addition, the Krebs cycle is the last metabolizing hub for the metabolism of lipids, carbohydrates, and amino acids and is associated with age-related and metabolic diseases [98, 101, 103]. For example, in the process of aspartate conversion to asparagine, ammonia is transferred from glutamine to asparagine which converts it to glutamate. In the Krebs cycle, glutamate is used as a substrate to produce α- ketoglutaraldehyde and subsequently succinate, critical intermediates of the Krebs cycle [104–106]. Glutamate production and consumption reach its highest level during an immune response in mammals [104–106]. It was found in a study that noninvasive 40 Hz flickering white light reduced Aβ load by recruiting microglia, immune cells in the brain, in AD mouse models [107]. Based on our observation, the flickering light might reduce Aβ plaque formation by triggering the immune response through activation of microglia in the brain of STZ-induced AD rats. Additionally, the 40 Hz light might induce glutamate metabolism and the Krebs cycle to improve energy production and normalize mitochondrial function [104–107].

The majority of ROS is produced by Complex I in the mitochondrial electron transport chain (ETC) [108]. Mitochondrial membrane integrity helps keep Complexes of ETC in close proximity, increase the efficiency of respiration, and reduce ROS production [109]. One possible explanation for elevated ROS in the AD rat brains in our findings could be a decrease in mitochondrial membrane integrity as evidenced by mitochondrial channel conductance and Complex I deficiency [110]. Another possibility is that the reverse electron transport (RET) mechanism is activated due to disruption in metabolite levels which leads to higher ROS production while keeping the ETC activity and mitochondrial function at minimal levels [111]. Studies have shown that high MMP, accumulated metabolic substrates, and reduced ubiquinone pools trigger this RET mechanism [111]. Our observation of high MMP and ROS associated with aberrant metabolic substrate levels concords with this proposed mechanism.

A series of amino acids including glutamate, glycine, and cysteine are involved in the process of reduced glutathione production which is the main ROS scavenger in the mitochondria [112]. Therefore, defects in the metabolism of these amino acids as indicated in our present study interfere with the renewal of ROS-scavenging function of glutathione. This, in turn, contributes to more build-up of ROS that might negatively impact ETC Complex activities and TCA cycle function [113]. Replenishing glutathione by re-activation of glutamatergic neurons and glutamate supply might explain the reducing effect of the 40 Hz flickering white light on ROS production [114]. The white light treatment might also facilitate the transmembrane transport of amino acids and metabolites required for mitochondrial membrane integrity, mitochondrial conductance and function [114].

The 40 Hz flickering white light has great therapeutic potential for treating neurodegenerative disorders since it has shown promising improvement in the pathophysiology of AD disease in animal models. Translational studies are needed to reproduce the outcome from animal models in humans. In addition, more animal studies are required to examine the duration of light therapy and changes in the plasma membrane BK_Ca_ channels in parallel with mitoBK_Ca_ channels to dissect the direct effect of light on each of these critical channels independently.

## Credit author statement

Amir Barzegar Behrooz (A.B.B.): Investigation, Visualization, Figure Illustration, Writing-Original Draft, Writing-Review & Editing. Friba Khodagholi (F.K.): Conceptualization, Methodology, Software, Validation, Writing- Review & Editing. Maryam Nazari (M.N.): Investigation, Formal analysis, Validation, Visualization. Hamid Latifi-Navid (H.L.N): Investigation, Formal analysis, Writing - Original Draft. Fatemeh vosogh (F.V.): Validation, Software. Mojdeh Anjomani (M.A.): Investigation, Validation, Software. Jaber Lotfi (J.L.): Investigation, Resources. M Ahmadiani (M.A.): Formal analysis, Validation. Afsaneh Eliassi (A.E.): Validation, Writing- Review & Editing. Fatemeh Nabavizadeh (F.N.): Resources, Validation. Elham soleimani: Formal analysis, Validation. Javad Fahanik-babaei (J.F.B): Conceptualization, Methodology, Software, Formal analysis, Validation, Supervision, Funding acquisition, Project administration, Writing- Review & Editing. All authors read and approved the final manuscript.

## Data availability statement

The original contributions presented in this study are included in the article/supplementary material; further inquiries can be directed to the corresponding author.

## Ethics statement

The Ethics and Research Committee of the Tehran University of Medical Sciences (IR TUMS.NI.REC.1398.056) reviewed and approved this animal study.

## Acknowledgments

We thank the Neuroscience Institute, Tehran University of Medical Sciences and Shahid Beheshti University of Medical Sciences for financial support.

## Supplementary material

There are 5 supplementary tables (tables 1-5) in excel files attached.

## References

1. Abubakar, M.B., et al., Alzheimer’s Disease: An Update and Insights Into Pathophysiology. Front Aging Neurosci, 2022. 14: p. 742408.

2. Knopman, D.S., et al., Alzheimer disease. Nat Rev Dis Primers, 2021. 7(1): p. 33.

3. Jack, C.R., Jr., et al., NIA-AA Research Framework: Toward a biological definition of Alzheimer’s disease. Alzheimers Dement, 2018. 14(4): p. 535–562.

4. Sperling, R.A., et al., Toward defining the preclinical stages of Alzheimer’s disease: recommendations from the National Institute on Aging-Alzheimer’s Association workgroups on diagnostic guidelines for Alzheimer’s disease. Alzheimers Dement, 2011. 7(3): p. 280–92.

5. Yiannopoulou, K.G. and S.G. Papageorgiou, Current and Future Treatments in Alzheimer Disease: An Update. J Cent Nerv Syst Dis, 2020. 12: p. 1179573520907397.

6. Chen, Y., A.K.Y. Fu, and N.Y. Ip, Synaptic dysfunction in Alzheimer’s disease: Mechanisms and therapeutic strategies. Pharmacol Ther, 2019. 195: p. 186–198.

7. Peters, A., C. Reisch, and D. Langemann, LTP or LTD? Modeling the Influence of Stress on Synaptic Plasticity. eNeuro, 2018. 5(1).

8. Johnstone, D.M., et al., Turning On Lights to Stop Neurodegeneration: The Potential of Near Infrared Light Therapy in Alzheimer’s and Parkinson’s Disease. Front Neurosci, 2015. 9: p. 500.

9. Hamblin, M.R., Shining light on the head: Photobiomodulation for brain disorders. BBA Clin, 2016. 6: p. 113–124.

10. Rojas, J.C. and F. Gonzalez-Lima, Neurological and psychological applications of transcranial lasers and LEDs. Biochem Pharmacol, 2013. 86(4): p. 447–57.

11. Iaccarino, H.F., et al., Gamma frequency entrainment attenuates amyloid load and modifies microglia. Nature, 2016. 540(7632): p. 230–235.

12. Adaikkan, C. and L.H. Tsai, Gamma Entrainment: Impact on Neurocircuits, Glia, and Therapeutic Opportunities. Trends Neurosci, 2020. 43(1): p. 24–41.

13. Moissenet, F., L. Modenese, and R. Dumas, Alterations of musculoskeletal models for a more accurate estimation of lower limb joint contact forces during normal gait: A systematic review. J Biomech, 2017. 63: p. 8–20.

14. Martorell, A.J., et al., Multi-sensory Gamma Stimulation Ameliorates Alzheimer’s-Associated Pathology and Improves Cognition. Cell, 2019. 177(2): p. 256–271.e22.

15. Saltmarche, A.E., et al., Significant Improvement in Cognition in Mild to Moderately Severe Dementia Cases Treated with Transcranial Plus Intranasal Photobiomodulation: Case Series Report. Photomed Laser Surg, 2017. 35(8): p. 432–441.

16. Nazari, M., et al., The Effect of 40-Hz White LED Therapy on Structure-Function of Brain Mitochondrial ATP-Sensitive Ca-Activated Large-Conductance Potassium Channel in Amyloid Beta Toxicity. Neurotox Res, 2022. 40(5): p. 1380–1392.

17. Adaikkan, C., et al., Gamma Entrainment Binds Higher-Order Brain Regions and Offers Neuroprotection. Neuron, 2019. 102(5): p. 929–943.e8.

18. Zheng, L., et al., Rhythmic light flicker rescues hippocampal low gamma and protects ischemic neurons by enhancing presynaptic plasticity. Nat Commun, 2020. 11(1): p. 3012.

19. Wang, W., et al., Gamma frequency entrainment rescues cognitive impairment by decreasing postsynaptic transmission after traumatic brain injury. CNS Neurosci Ther, 2023. 29(4): p. 1142–1153.

20. Chan, D., et al., Gamma frequency sensory stimulation in mild probable Alzheimer’s dementia patients: Results of feasibility and pilot studies. PLoS One, 2022. 17(12): p. e0278412.

21. Manczak, M., et al., Mitochondria are a direct site of A beta accumulation in Alzheimer’s disease neurons: implications for free radical generation and oxidative damage in disease progression. Hum Mol Genet, 2006. 15(9): p. 1437–49.

22. Sanz-Blasco, S., et al., Mitochondrial Ca2+ overload underlies Abeta oligomers neurotoxicity providing an unexpected mechanism of neuroprotection by NSAIDs. PLoS One, 2008. 3(7): p. e2718.

23. Fisher-Wellman, K.H., et al., Pyruvate dehydrogenase complex and nicotinamide nucleotide transhydrogenase constitute an energy-consuming redox circuit. Biochem J, 2015. 467(2): p. 271–80.

24. Müller, W.E., et al., Mitochondrial dysfunction: common final pathway in brain aging and Alzheimer’s disease--therapeutic aspects. Mol Neurobiol, 2010. 41(2-3): p. 159–71.

25. Mutisya, E.M., A.C. Bowling, and M.F. Beal, Cortical cytochrome oxidase activity is reduced in Alzheimer’s disease. J Neurochem, 1994. 63(6): p. 2179–84.

26. Hamblin, M.R., Photobiomodulation for Alzheimer’s Disease: Has the Light Dawned? Photonics, 2019. 6(3).

27. Lu, Y., et al., Low-level laser therapy for beta amyloid toxicity in rat hippocampus. Neurobiol Aging, 2017. 49: p. 165–182.

28. Nazari, M., et al., The 40-Hz White Light-Emitting Diode (LED) Improves the Structure– Function of the Brain Mitochondrial KATP Channel and Respiratory Chain Activities in Amyloid Beta Toxicity. Molecular Neurobiology, 2022. 59(4): p. 2424–2440.

29. Nazari, M., et al., The Effect of 40-Hz White LED Therapy on Structure–Function of Brain Mitochondrial ATP-Sensitive Ca-Activated Large-Conductance Potassium Channel in Amyloid Beta Toxicity. Neurotoxicity Research, 2022. 40(5): p. 1380–1392.

30. Garcia-Gil, M., et al., Metabolic Aspects of Adenosine Functions in the Brain. Front Pharmacol, 2021. 12: p. 672182.

31. Borea, P.A., et al., Pharmacology of Adenosine Receptors: The State of the Art. Physiol Rev, 2018. 98(3): p. 1591–1625.

32. Cunha, R.A., How does adenosine control neuronal dysfunction and neurodegeneration? J Neurochem, 2016. 139(6): p. 1019–1055.

33. Chen, J.F., H.K. Eltzschig, and B.B. Fredholm, Adenosine receptors as drug targets-- what are the challenges? Nat Rev Drug Discov, 2013. 12(4): p. 265–86.

34. Szabo, I. and A. Szewczyk, Mitochondrial Ion Channels. Annual Review of Biophysics, 2023. 52(1): p. 229–254.

35. Trombetta-Lima, M., I.E. Krabbendam, and A.M. Dolga, Calcium-activated potassium channels: implications for aging and age-related neurodegeneration. Int J Biochem Cell Biol, 2020. 123: p. 105748.

36. Fahanik-Babaei, J., et al., Electro-pharmacological profile of a mitochondrial inner membrane big-potassium channel from rat brain. Biochim Biophys Acta, 2011. 1808(1): p. 454–60.

37. Fahanik-Babaei, J., A. Eliassi, and R. Saghiri, How many types of large conductance CaLJ²-activated potassium channels exist in brain mitochondrial inner membrane: evidence for a new mitochondrial large conductance Ca²LJ-activated potassium channel in brain mitochondria. Neuroscience, 2011. 199: p. 125–32.

38. Jafari, A., et al., Brain mitochondrial ATP-insensitive large conductance Ca(+)(2)- activated K(+) channel properties are altered in a rat model of amyloid-beta neurotoxicity. Exp Neurol, 2015. 269: p. 8–16.

39. Torabi, N., et al., Intranasal insulin improves the structure-function of the brain mitochondrial ATP-sensitive Ca(2+) activated potassium channel and respiratory chain activities under diabetic conditions. Biochim Biophys Acta Mol Basis Dis, 2021. 1867(4): p. 166075.

40. Vorhees, C.V. and M.T. Williams, Morris water maze: procedures for assessing spatial and related forms of learning and memory. Nat Protoc, 2006. 1(2): p. 848–58.

41. Antunes, M. and G. Biala, The novel object recognition memory: neurobiology, test procedure, and its modifications. Cognitive Processing, 2012. 13(2): p. 93–110.

42. Gupta, P., et al., Intracerebroventricular Abeta-Induced Neuroinflammation Alters Peripheral Immune Responses in Rats. J Mol Neurosci, 2018. 66(4): p. 572–586.

43. Navarro, A., et al., Vitamin E at high doses improves survival, neurological performance, and brain mitochondrial function in aging male mice. Am J Physiol Regul Integr Comp Physiol, 2005. 289(5): p. R1392–9.

44. Rosenthal, R.E., et al., Cerebral ischemia and reperfusion: prevention of brain mitochondrial injury by lidoflazine. J Cereb Blood Flow Metab, 1987. 7(6): p. 752–8.

45. Fahanik-Babaei, J., et al., Electro-pharmacological profiles of two brain mitoplast anion channels: Inferences from single channel recording. Excli j, 2017. 16: p. 531–545.

46. Da Cruz, S., et al., Proteomic analysis of the mouse liver mitochondrial inner membrane. J Biol Chem, 2003. 278(42): p. 41566–71.

47. Singleton, W.S., M.S. Gray, M.L. Brown, and J.L. White, CHROMATOGRAPHICALLY HOMOGENEOUS LECITHIN FROM EGG PHOSPHOLIPIDS. J Am Oil Chem Soc, 1965. 42: p. 53–6.

48. Nabavi Zadeh, F., et al., Pre- and post-treatment of α-Tocopherol on cognitive, synaptic plasticity, and mitochondrial disorders of the hippocampus in icv-streptozotocin-induced sporadic Alzheimer’s-like disease in male Wistar rat. Front Neurosci, 2023. 17: p. 1073369.

49. Rahmani, P., et al., Determination of carnitine ester profile in the children with type 1 diabetes: a valuable step towards a better management. Arch Physiol Biochem, 2022. 128(5): p. 1209–1214.

50. Spinazzi, M., et al., Assessment of mitochondrial respiratory chain enzymatic activities on tissues and cultured cells. Nat Protoc, 2012. 7(6): p. 1235–46.

51. Luo, J. and R. Shi, Acrolein induces oxidative stress in brain mitochondria. Neurochem Int, 2005. 46(3): p. 243–52.

52. Pipatpiboon, N., W. Pratchayasakul, N. Chattipakorn, and S.C. Chattipakorn, PPARγ agonist improves neuronal insulin receptor function in hippocampus and brain mitochondria function in rats with insulin resistance induced by long term high-fat diets. Endocrinology, 2012. 153(1): p. 329–38.

53. Pang, Z., et al., MetaboAnalyst 6.0: towards a unified platform for metabolomics data processing, analysis and interpretation. Nucleic Acids Res, 2024.

54. Luo, W., et al., Pathview Web: user friendly pathway visualization and data integration. Nucleic Acids Res, 2017. 45(W1): p. W501–W508.

55. Fahanik-Babaei, J., A. Eliassi, and R. Saghiri, How many types of large conductance Ca+2-activated potassium channels exist in brain mitochondrial inner membrane: evidence for a new mitochondrial large conductance Ca2+-activated potassium channel in brain mitochondria. Neuroscience, 2011. 199: p. 125–132.

56. Hosseininia, M., et al., Memory impairment was ameliorated by corticolimbic microinjections of arachidonylcyclopropylamide (ACPA) and miRNA-regulated lentiviral particles in a streptozotocin-induced Alzheimer’s rat model. Exp Neurol, 2023. 370: p. 114560.

57. Adeli, S., M. Zahmatkesh, and M. Ansari Dezfouli, Simvastatin Attenuates Hippocampal MMP-9 Expression in the Streptozotocin-Induced Cognitive Impairment. Iran Biomed J, 2019. 23(4): p. 262–71.

58. Khuman, J., et al., Low-level laser light therapy improves cognitive deficits and inhibits microglial activation after controlled cortical impact in mice. J Neurotrauma, 2012. 29(2): p. 408–17.

59. Pan, W.T., P.M. Liu, D. Ma, and J.J. Yang, Advances in photobiomodulation for cognitive improvement by near-infrared derived multiple strategies. J Transl Med, 2023. 21(1): p. 135.

60. Semyachkina-Glushkovskaya, O., et al., Night Photostimulation of Clearance of Beta-Amyloid from Mouse Brain: New Strategies in Preventing Alzheimer’s Disease. Cells, 2021. 10(12).

61. Park, S.S., et al., Combined effects of aerobic exercise and 40-Hz light flicker exposure on early cognitive impairments in Alzheimer’s disease of 3×Tg mice. J Appl Physiol (1985), 2022. 132(4): p. 1054–1068.

62. Nobili, A., et al., Dopamine neuronal loss contributes to memory and reward dysfunction in a model of Alzheimer’s disease. Nat Commun, 2017. 8: p. 14727.

63. Storga, D., K. Vrecko, J.G. Birkmayer, and G. Reibnegger, Monoaminergic neurotransmitters, their precursors and metabolites in brains of Alzheimer patients. Neurosci Lett, 1996. 203(1): p. 29–32.

64. Gloria, Y., et al., Dopaminergic dysfunction in the 3xTg-AD mice model of Alzheimer’s disease. Sci Rep, 2021. 11(1): p. 19412.

65. Babić Leko, M., P.R. Hof, and G. Šimić, Chapter 9 - Alterations and interactions of subcortical modulatory systems in Alzheimer’s disease, in Progress in Brain Research, G. Di Giovanni and P. De Deurwaerdere, Editors. 2021, Elsevier. p. 379–421.

66. Harrison, S.J., et al., Light therapy and serotonin transporter binding in the anterior cingulate and prefrontal cortex. Acta Psychiatr Scand, 2015. 132(5): p. 379–88.

67. Tyrer, A.E., et al., Serotonin transporter binding is reduced in seasonal affective disorder following light therapy. Acta Psychiatr Scand, 2016. 134(5): p. 410–419.

68. Tomaz de Magalhaes, M., S.C. Nunez, I.T. Kato, and M.S. Ribeiro, Light therapy modulates serotonin levels and blood flow in women with headache. A preliminary study. Exp Biol Med (Maywood), 2016. 241(1): p. 40–5.

69. Reppert, S.M., et al., Molecular characterization of a second melatonin receptor expressed in human retina and brain: the Mel1b melatonin receptor. Proc Natl Acad Sci U S A, 1995. 92(19): p. 8734–8.

70. Dubocovich, M.L., Melatonin is a potent modulator of dopamine release in the retina. Nature, 1983. 306(5945): p. 782–4.

71. Tian, J., E. Du, and L. Guo, Mitochondrial Interaction with Serotonin in Neurobiology and Its Implication in Alzheimer’s Disease. J Alzheimers Dis Rep, 2023. 7(1): p. 1165–1177.

72. Harris, J.J., R. Jolivet, and D. Attwell, Synaptic energy use and supply. Neuron, 2012. 75(5): p. 762–77.

73. Skaper, S.D., L. Facci, M. Zusso, and P. Giusti, Synaptic Plasticity, Dementia and Alzheimer Disease. CNS Neurol Disord Drug Targets, 2017. 16(3): p. 220–233.

74. Bliss, T.V. and T. Lomo, Long-lasting potentiation of synaptic transmission in the dentate area of the anaesthetized rabbit following stimulation of the perforant path. J Physiol, 1973. 232(2): p. 331–56.

75. Rai, S., P.K. Kamat, C. Nath, and R. Shukla, Glial activation and post-synaptic neurotoxicity: the key events in Streptozotocin (ICV) induced memory impairment in rats. Pharmacol Biochem Behav, 2014. 117: p. 104–17.

76. Catterall, W.A., K. Leal, and E. Nanou, Calcium Channels and Short-term Synaptic Plasticity*. Journal of Biological Chemistry, 2013. 288(15): p. 10742–10749.

77. Leszkiewicz, D.N., K. Kandler, and E. Aizenman, Enhancement of NMDA receptor-mediated currents by light in rat neurones in vitro. J Physiol, 2000. 524 **Pt 2**(Pt 2): p. 365–74.

78. Wang, W., et al., Mitochondria dysfunction in the pathogenesis of Alzheimer’s disease: recent advances. Mol Neurodegener, 2020. 15(1): p. 30.

79. Rossi, A., P. Pizzo, and R. Filadi, Calcium, mitochondria and cell metabolism: A functional triangle in bioenergetics. Biochim Biophys Acta Mol Cell Res, 2019. 1866(7): p. 1068–1078.

80. Torabi, N., et al., Intranasal insulin improves the structure–function of the brain mitochondrial ATP–sensitive Ca2+ activated potassium channel and respiratory chain activities under diabetic conditions. Biochimica et Biophysica Acta (BBA) - Molecular Basis of Disease, 2021. 1867(4): p. 166075.

81. Brüning, J.C., et al., Role of brain insulin receptor in control of body weight and reproduction. Science, 2000. 289(5487): p. 2122–5.

82. Kullmann, S., et al., Brain Insulin Resistance at the Crossroads of Metabolic and Cognitive Disorders in Humans. Physiol Rev, 2016. 96(4): p. 1169–209.

83. Arnold, S.E., et al., Brain insulin resistance in type 2 diabetes and Alzheimer disease: concepts and conundrums. Nat Rev Neurol, 2018. 14(3): p. 168–181.

84. O’Malley, D., L.J. Shanley, and J. Harvey, Insulin inhibits rat hippocampal neurones via activation of ATP-sensitive K+ and large conductance Ca2+-activated K+ channels. Neuropharmacology, 2003. 44(7): p. 855–63.

85. Krabbendam, I.E., B. Honrath, C. Culmsee, and A.M. Dolga, Mitochondrial Ca(2+)- activated K(+) channels and their role in cell life and death pathways. Cell Calcium, 2018. 69: p. 101–111.

86. Palazzo, E., F. Rossi, V. de Novellis, and S. Maione, Endogenous modulators of TRP channels. Curr Top Med Chem, 2013. 13(3): p. 398–407.

87. Gorriz, R.F. and P. Imhof, Interplay of Hydration and Protonation Dynamics in the K- Channel of Cytochrome c Oxidase. Biomolecules, 2022. 12(11).

88. Kozyreva, T.V., I.V. Orlov, A.R. Boyarskaya, and I.P. Voronova, Hypothalamic TRPM8 and TRPA1 ion channel genes in the regulation of temperature homeostasis at water balance changes. Neurosci Lett, 2024. 828: p. 137763.

89. Yang, L., H. Youngblood, C. Wu, and Q. Zhang, Mitochondria as a target for neuroprotection: role of methylene blue and photobiomodulation. Transl Neurodegener, 2020. 9(1): p. 19.

90. Wang, H., et al., Recent Advances in Chemical Biology of Mitochondria Targeting. Front Chem, 2021. 9: p. 683220.

91. Bednarczyk, P., A. Koziel, W. Jarmuszkiewicz, and A. Szewczyk, Large-conductance Ca²LJ-activated potassium channel in mitochondria of endothelial EA.hy926 cells. Am J Physiol Heart Circ Physiol, 2013. 304(11): p. H1415–27.

92. Ball, K.A., P.R. Castello, and R.O. Poyton, Low intensity light stimulates nitrite-dependent nitric oxide synthesis but not oxygen consumption by cytochrome c oxidase: Implications for phototherapy. J Photochem Photobiol B, 2011. 102(3): p. 182–91.

93. Waypa, G.B., K.A. Smith, and P.T. Schumacker, O2 sensing, mitochondria and ROS signaling: The fog is lifting. Mol Aspects Med, 2016. 47-48: p. 76–89.

94. Kulawiak, B., A.P. Kudin, A. Szewczyk, and W.S. Kunz, BK channel openers inhibit ROS production of isolated rat brain mitochondria. Exp Neurol, 2008. 212(2): p. 543–7.

95. Wang, L., et al., Cognitive recovery by chronic activation of the large-conductance calcium-activated potassium channel in a mouse model of Alzheimer’s disease. Neuropharmacology, 2015. 92: p. 8–15.

96. Feng, B., S. Raghavachari, and J. Lisman, Quantitative estimates of the cytoplasmic, PSD, and NMDAR-bound pools of CaMKII in dendritic spines. Brain Res, 2011. 1419: p. 46–52.

97. Sibarov, D.A. and S.M. Antonov, Calcium-Dependent Desensitization of NMDA Receptors. Biochemistry (Mosc), 2018. 83(10): p. 1173–1183.

98. Chen, M., et al., Investigation into potential mechanisms of metabolic syndrome by integrative analysis of metabolomics and proteomics. PLoS One, 2022. 17(7): p. e0270593.

99. van Gelderen, T.A., et al., Metabolomic and transcriptomic profiles after immune stimulation in the zebrafish testes. Genomics, 2023. 115(2): p. 110581.

100. Robinson, J.B., Jr. and P.A. Srere, Organization of Krebs tricarboxylic acid cycle enzymes in mitochondria. J Biol Chem, 1985. 260(19): p. 10800–5.

101. Villa, R.F., A. Gorini, and S. Hoyer, Effect of ageing and ischemia on enzymatic activities linked to Krebs’ cycle, electron transfer chain, glutamate and aminoacids metabolism of free and intrasynaptic mitochondria of cerebral cortex. Neurochem Res, 2009. 34(12): p. 2102–16.

102. Aghanoori, M.R., et al., Sensory neurons derived from diabetic rats exhibit deficits in functional glycolysis and ATP that are ameliorated by IGF-1. Mol Metab, 2021. 49: p. 101191.

103. Pinkosky, S.L., P.H.E. Groot, N.D. Lalwani, and G.R. Steinberg, Targeting ATP-Citrate Lyase in Hyperlipidemia and Metabolic Disorders. Trends Mol Med, 2017. 23(11): p. 1047–1063.

104. Lomelino, C.L., J.T. Andring, R. McKenna, and M.S. Kilberg, Asparagine synthetase: Function, structure, and role in disease. J Biol Chem, 2017. 292(49): p. 19952–19958.

105. Newsholme, P., V.L.S. Diniz, G.T. Dodd, and V. Cruzat, Glutamine metabolism and optimal immune and CNS function. Proc Nutr Soc, 2023. 82(1): p. 22–31.

106. Choi, I., H. Son, and J.H. Baek, Tricarboxylic Acid (TCA) Cycle Intermediates: Regulators of Immune Responses. Life (Basel), 2021. 11(1).

107. Singer, A.C., et al., Noninvasive 40-Hz light flicker to recruit microglia and reduce amyloid beta load. Nat Protoc, 2018. 13(8): p. 1850–1868.

108. Kushnareva, Y., A.N. Murphy, and A. Andreyev, Complex I-mediated reactive oxygen species generation: modulation by cytochrome c and NAD(P)+ oxidation-reduction state. Biochem J, 2002. 368(Pt 2): p. 545–53.

109. Novack, G.V., P. Galeano, E.M. Castano, and L. Morelli, Mitochondrial Supercomplexes: Physiological Organization and Dysregulation in Age-Related Neurodegenerative Disorders. Front Endocrinol (Lausanne), 2020. 11: p. 600.

110. Sharma, L.K., et al., Mitochondrial respiratory complex I dysfunction promotes tumorigenesis through ROS alteration and AKT activation. Hum Mol Genet, 2011. 20(23): p. 4605–16.

111. Onukwufor, J.O., B.J. Berry, and A.P. Wojtovich, Physiologic Implications of Reactive Oxygen Species Production by Mitochondrial Complex I Reverse Electron Transport. Antioxidants (Basel), 2019. 8(8).

112. Desideri, E., F. Ciccarone, and M.R. Ciriolo, Targeting Glutathione Metabolism: Partner in Crime in Anticancer Therapy. Nutrients, 2019. 11(8).

113. Forrester, S.J., et al., Reactive Oxygen Species in Metabolic and Inflammatory Signaling. Circ Res, 2018. 122(6): p. 877–902.

114. Zhou, X., et al., 40 Hz light flickering promotes sleep through cortical adenosine signaling. Cell Res, 2024. 34(3): p. 214–231.

